# Evidence of active oviposition avoidance to systemically applied imidacloprid in the Colorado Potato Beetle (*Leptinotarsa decemlineata*)

**DOI:** 10.1101/2022.07.11.499582

**Authors:** Alitha Edison, Anja Michelbach, Dominique Sowade, Luise Schmidt, Martin Schäfer, Ralf Nauen, Pablo Duchen, Shuqing Xu

## Abstract

Agricultural pests can develop behavioral resistance to insecticides via choosing to feed or oviposit on non-toxic hosts. As young larvae have relatively low mobility, oviposition preferences from female adults may play a critical role in shaping the evolutionary trajectory of pest populations. While oviposition avoidance of toxic hosts was found in different agriculture pests, it remains unclear whether such preferences can be learned from female adults. To address this question, we investigated feeding and oviposition preferences to imidacloprid in the Colorado Potato Beetle (CPB, *Leptinotarsa decemlineata*), a major potato pest. We first identified two CPB strains that have different levels of resistance to imidacloprid. Then, we performed choice assays in the two strains and found that both strains did not have an innate feeding avoidance to systemically applied imidacloprid at both larval and adult stages. Further oviposition choice assays showed that the susceptible strain preferred to lay eggs on insecticide-free plants while the resistant strain did not. Analysing moving patterns of the two strains suggested that the oviposition preference is likely due to active learning by the female adults. Together, these results indicate that CPB can have active oviposition avoidance, which might have contributed to the rapid global invasion of this agricultural pest.

## 1. Introduction

Rapid evolution of insecticide resistance threatens both global food supply and human health (1). Managing the evolution of insecticide resistance in pests requires integrated investigations on the underlying physiological, biochemical, and behavioral mechanisms. While significant progress has been made in identifying physiological and biochemical mechanisms (2–4), the extent to which behavior contributes to rapid evolution of insecticide resistance remains largely unclear. For instance, insects can use avoidance behaviors to feed or live on non-toxic hosts over toxic hosts and thus survive insecticides. Selection could therefore theoretically favor such behavioral traits (5–7). Because young larvae have limited ability to move and are more susceptible to food quality, oviposition choice from adult female plays a critical role in determining the fitness of larvae. As many adult insects feed on the host before oviposition, it is possible that female adults can detect the toxicity of hosts and actively choose to lay eggs in non-toxic hosts. However, this has never been experimentally verified before, despite the existence of oviposition avoidance behavior in several insects (8– 11).

The Colorado Potato Beetle (*Leptinotarsa decemlineata*) is a global pest that can cause complete defoliation of potato plants (12). Since expanding its hosts to potato about 200 years ago in Mexico, where it is native, the beetle has rapidly spread to North America, Europe, and now reached Asia (13). Currently, CPB is known to be resistant to over 54 different insecticides including the neonicotinoid imidacloprid, commonly used in agriculture (14). Increased metabolism, target site insensitivity, reduced insecticide penetration, and enhanced excretion are some of the known mechanisms of resistance in CPB (15). While some studies did observe behavioral differences between resistant and susceptible CPB strains (16,17), their experimental designs did not allow disentanglement of the physiological effects of toxin consumption from the behavioral resistance (17). Thus, to which extent behavior contributes to the evolution of insecticide resistance in CPB remains largely unknown.

CPB often prefers domesticated potato (*Solanum tuberosum*) plants as their hosts, both for feeding and oviposition, although some contradictory evidence from different geographical populations exist (18–21). Host finding is facilitated by visual and olfactory cues from a distance (22–25). Once they are within a close range, the larvae and beetles are attracted to volatiles associated with *Solanum* plants (23,26). After finding a host, specific orientation and a series of sequential feeding behaviors that follow upon contact determine its acceptance or rejection (22,27–29). During their elaborate biting and sampling routine, CPB adults and larvae use their sensitive gustatory cells to taste the compounds in a given host to determine whether it is suitable (29–32). Experience or learning influences the host preference in CPB (33) as it does in many insects (34–36).

Feeding preferences by the larvae and adults are generally similar, although the host plant range of the larvae may be more restricted (37). Once a host is accepted, continued feeding, mating and oviposition subsequently follow. Oviposition preference is generally thought to be consistent with the feeding preference because the gravid females rarely succeed in flight, but there is some evidence for discrepancies between feeding and oviposition preferences (19,21). However, all of feeding and oviposition preference of CPB were measured among different plant species. To our knowledge, no study has examined the feeding and oviposition preference between insecticide treated and untreated plants in CPB.

Here, we aimed to investigate CPB’s behavioral response to systemically applied imidacloprid, both in terms of feeding and oviposition choices. To this end, we first screened five different CPB strains and identified a susceptible and a moderately resistant strain. Then, we quantified the feeding and oviposition preferences of these two strains and aimed to address the two main questions: 1) Do CPB larvae and/or adults actively avoid feeding on insecticide-treated plants? 2) Do adult females actively avoid insecticide-treated plants for oviposition? Answers to these questions will sharpen our understanding of behavioral resistance in a major agricultural pest, which might aid the successful implementation of integrated pest management strategies in the future.

## 2. Materials & Methods

### A. Plants, Insects & Insecticide

Five-week-old *Solanum tuberosum* plants (Annabelle variety, purchased from TOLLS Kartoffelhandel GmbH & Co. KG, Willich) were used for rearing the beetles, and for the experiments. The plants were grown in trays of size 39cm x 29cm x 7cm in a greenhouse under a long day photoperiod (16h light: 8h dark), and a temperature of 24°C. The rearing of the beetles and experiments were all carried out in the same greenhouse, also under a long day photoperiod and a temperature of 24°C.

Colorado potato beetles were supplied by Dr. Ralf Nauen, Bayer AG (Monheim, Germany). The five strains used in the study were collected by the supplier and group from different parts in Europe at different points in time (38). They are namely, D01 (Germany, 2002), E01 (Spain, 2014), E02 (Spain, 2017), E06 (Spain, 2012), and U01 (Ukraine, 2012). All insects had been reared under the same conditions (without the addition of insecticides) for at least 10 generations in 85cm x 45cm x 55cm sized insect cages. During each generation, eggs were collected regularly using wet brushes and stored at 10°C for synchronizing hatching times. Afterwards they were moved back into the greenhouse. Water was sprayed on top of the eggs to prevent drying. After five days, when the larvae hatched, they were introduced into cages with sufficient potato plants where they were allowed to feed on the leaves and develop. The watering of the plants was stopped once the larvae reached the final stage of development, which was implied by the larger size and the colour change from red to orange. Then, the larvae were allowed to fall into the trays and pupate within the soil. When the adults emerged 11 days later, they were supplied with fresh potato plants.

The neonicotinoid insecticide, imidacloprid was used in our experiments in the form of water dispersible granules available commercially by the name Confidor WG 70 (70g/Kg Imidacloprid, Bayer AG, Monheim, Germany). The insecticide granules were mixed in water and applied to the soil.

*Insecticide quantification* was performed to monitor the amount of imidacloprid in the leaves of potato plants after watering their soil with an aqueous solution of 2ppm imidacloprid (insecticide-treated) or with pure water (control). The experiments were conducted with five-week-old plants and five replicates were collected 0, 1, 2, 3, 5, 7, 10 and 14 days following the treatment. Samples were collected by cutting leaf discs with a 1.75cm diameter cork borer from the leaves at the 3^rd^, 4^th^ or 5^th^ position (from the top) of insecticide-treated and control plants. The leaf discs were snap frozen in liquid nitrogen and thoroughly homogenized using a Retsch ball mill. Approximately 10-15 mg (fresh weight or FW) of finely ground plant material was extracted with 1ml of 70% MeOH with 1% formic acid (Fisher Chemical) as extraction buffer supplemented with 10ng/ml D_4_-Imidacloprid (Sigma-Aldrich) as an internal standard. After thoroughly shaking, the samples were incubated for 30min on ice, centrifuged at 16900g for 20min at 4°C and subsequently the supernatant was collected. The pellet was re-extracted with 0.5ml extraction buffer without internal standard, by another round of shaking, centrifugation and collecting the supernatant. Both supernatants were combined and centrifuged again to remove remaining particles before analysis.

Chromatography was performed on a Nexera X3 UHPLC system (Shimadzu). For separation a Zorbax RRHD Eclipse XDB C18 column (1.8µm, 3×50mm; Agilent) was used. The mobile phase comprised solvent A (water, 0.1% (v/v) formic acid, 0.05% (v/v) acetonitrile) and solvent B (methanol) with following elution profile: 0min, 10% B; 0.5min, 10% B; 1min, 25% B; 4min, 30% B; 4.5min, 100% B; 5.5min, 100% B; 6min, 10% B; 7min, 10% B; with a flow rate of 0.5mL/min. The column temperature was maintained at 42°C. The chromatography was coupled to a LCMS-8060 mass spectrometer (Shimadzu) with an ESI source. The instrument was operated in positive ionization mode multi-reaction-monitoring modus to analyze the parent ion → product ion transitions of the analytes (Supplementary Table. 1). Following source parameters were used: nebulizing gas flow, 3 L/min; heating gas flow,10 L/min; drying gas flow, 10 L/min; interface temperature, 300°C; DL temperature, 250°C; heat block temperature, 400°C. The CID gas flow was set to 270kPa and the resolution for Q1 and Q3 to unit resolution.

Based on the determined insecticide levels, for all subsequent experiments, the plants were treated two days before the start of the experiment to ensure enough insecticide had reached the leaves (Supplementary Fig. 1).

### B. Toxicity assays with larvae

Preliminary information on the concentrations was first obtained from previous studies (39–42). However, the values highly varied depending on the geographical area and time. Based on the manufacturer’s recommendation to use 245 ppm for field application, we speculated this concentration would cause 100% mortality. However, 100% mortality was not observed for all the strains with the concentration of 245 ppm. Because of this, more pretests were done with 500ppm and 1000ppm insecticide solutions until the concentration that caused close to 100% mortality for all strains was found (1000ppm). Once this value was obtained, three points in between 0 and 1000 were chosen to facilitate plotting the dose-response curves.

Toxicity assays were performed with two-day-old larvae using two different methods: the topical application method, and the intact plants method. Because the first method is a standard test recommended by the Insecticide Resistance Action Committee (43) for leaf eating insects, it has the advantage of producing results that are comparable with other studies that use the same method. On the other hand, the method using intact plants more closely resembles a natural setting. Another important feature to note here is that in the first method, the larvae are forced to take up the insecticide, although the variation in cuticle thickness might play a deciding role. Meanwhile in the second method, the larvae can quit or slow down feeding to recover from the stress caused by ingestion of the insecticide.

For topical applications, two-day-old larvae were put on a petri dish lined with filter paper, following the IRAC Susceptibility Test Method 029 (44). For each of the five strains (D01, E01, E02, E06, U01), five concentrations of insecticide (15.62ppm, 62.5ppm, 250ppm, 1000ppm) and a control (water without insecticide) were tested. For each concentration, six replicates were tested, and each replicate contained five larvae. We applied 1µl of the solution for the respective treatment (control or one of the 5 insecticide solutions) on the top of each larva. The larvae were supplied with enough leaves to feed on throughout the duration of the assay. Mortality was recorded after 48h.

For applying the insecticide via plants feeding, two-day-old larvae were placed on the middle leaves (positions 3-5 from the top) of plants treated with insecticide via soil application two days prior. For each of the five strains, four concentrations of insecticide (15.31ppm, 30.62ppm, 61.25ppm, 122.50ppm) and a control (water without insecticide) were tested. The concentrations were determined based on similar pretests as described above. For each concentration, two replicates, each containing 20 larvae, were used. Mortality was recorded after 48h.

### C. Choice assays (Feeding/Oviposition)

Based on the toxicity assay experiments (section B), a susceptible (D01) and a resistant strain (E06) were selected for the choice assays.

*Behavior assays with larvae* were performed in a petri dish setup. The innate feeding preference, that is, a preference that preceded any experience with a host, was tested using freshly hatched naïve larvae. To assess the effects of pre-feeding, larvae that had been fed with control or insecticide-treated plants were used. For each assay, the preference of 60 individuals was tested for the D01 and E06 strain, each. For each individual choice test, a single larva was placed in a petri dish arena with two choices of leaf discs. The leaf discs were cut using cork-borers of 1.75cm diameter. The insecticide-containing (T) leaf discs were cut from plants that had been watered with 1L of an aqueous insecticide solution containing 2ppm imidacloprid two days before the experiment. The insecticide-free leaf discs were cut from plants that had been treated with 1L of a control solution (water without insecticide) two days before the experiment. The petri dishes were lined with a filter paper and lightly sprayed with water to slow down drying up of the leaf discs. The two types of leaf discs were set up on opposite sides. The larva was picked up with a wet paintbrush and left at the center of the petri dish arena. During 4h of continuous observation, first choices were noted.

*Behavior assays with adults* were performed using a cage setup. The adults used in the experiment had freshly emerged from the soil. Their development during the larval stage was on insecticide-free plants. The feeding and oviposition choice behavior were closely monitored for adults of both of the strains in a cage setup in which both insecticide-free (C) and insecticide-treated (T) plants were present. The insecticide-treated plants were treated with 1L of an aqueous insecticide solution containing 2ppm imidacloprid two days before the experiment. The insecticide-free plants were treated with 1L of a control solution (water without insecticide) two days before the experiment. Eight male and eight female adults were introduced into each cage with a 15min interval between the sexes (order of introduction randomized between replicates). For each strain, 10 replicates were used. The feeding behavior of the adults was closely observed for 30 min and the first feeding choice (either C or T plants) of each individual was recorded for both sexes. In the following days, the number of beetles in each of the trays was recorded twice daily to track the beetle movement patterns. The eggs were collected daily to track the oviposition preferences. In addition, the number of individual clusters were also recorded. After 10 days, the total number of eggs and the number of egg clusters were calculated. The net mortality and percentage leaf damage were recorded at the end of the experiment after 10 days, based on counts and visual inspection respectively.

### D. Data analysis

All analyses were done using the “stats” package (and functions therein) implemented in R Core Team (2020). For estimating LC50 and LC90, the lowest concentrations of imidacloprid in the insecticide-water solution required to kill 50% and 90%, respectively, a generalized linear model (GLM) assuming a binomial family was used. The standard error were calculated using the dose.p() function in the MASS package(45).

The choice preference was quantified using the choice index (*CI*), which is defined as: 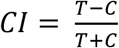, where *T* and *C* represent the ‘choice’ for insecticide-treated plants and control plants, respectively. ‘Choice’ is used here as a general term to denote the number of larvae, the number of beetles or the number of eggs. This index gives values above 0 if *T* is preferred and values below 0 if *C* is preferred. To analyze the feeding choice of the larvae, GLM were fitted to the data assuming a binomial distribution. The response variable, choice index was modelled as a function of strain and pre-feeding treatment. To analyze the first feeding choice of the adults, GLM were fitted to the data assuming a Poisson distribution. The response variable, choice index was modelled as a function of strain. To analyze the effect of sex, another GLM was fitted with the choice index as a function of strain and sex.

For the leaf damage, total number of eggs and total number of egg clusters data, GLMs were fitted assuming a negative binomial distribution. The response variables corresponding to these data, percentage leaf damage, total number of eggs and total number of egg clusters, were each modelled as a function of strain and treatment. For the beetle movement pattern and oviposition pattern data, the choice index was modelled using Linear Mixed Models with the strain as a fixed effect and the day as a random effect. P-values were determined using analysis of variance (ANOVA).

## 3. Results

### A. Toxicity assays using 1^st^ instar larvae

To determine the levels of susceptibility of the five different strains of CPB, we performed dose-response experiments using the first instar larvae (see Methods), both via topical applications and via plant feedings. For the intact plants method, among all five strains, E06 strain is the most resistant one, which is almost three times more resistant to imidacloprid than the most susceptible strain (D01) (Figure 1A). The E06 strain had the highest LC50 in the topical application method as well, but there are differences in other strains between the two methods, possibly due to differences in cuticle thickness, detoxification processes or behavioral differences (Figure 1B). The lowest average concentrations to kill 50% (LC50) and 90% (LC90) of the larvae using the intact plants method are shown for all five strains in Table 1. LC50 and LC90 values for the topical application method are shown in Table 2.

**Figure 1.**
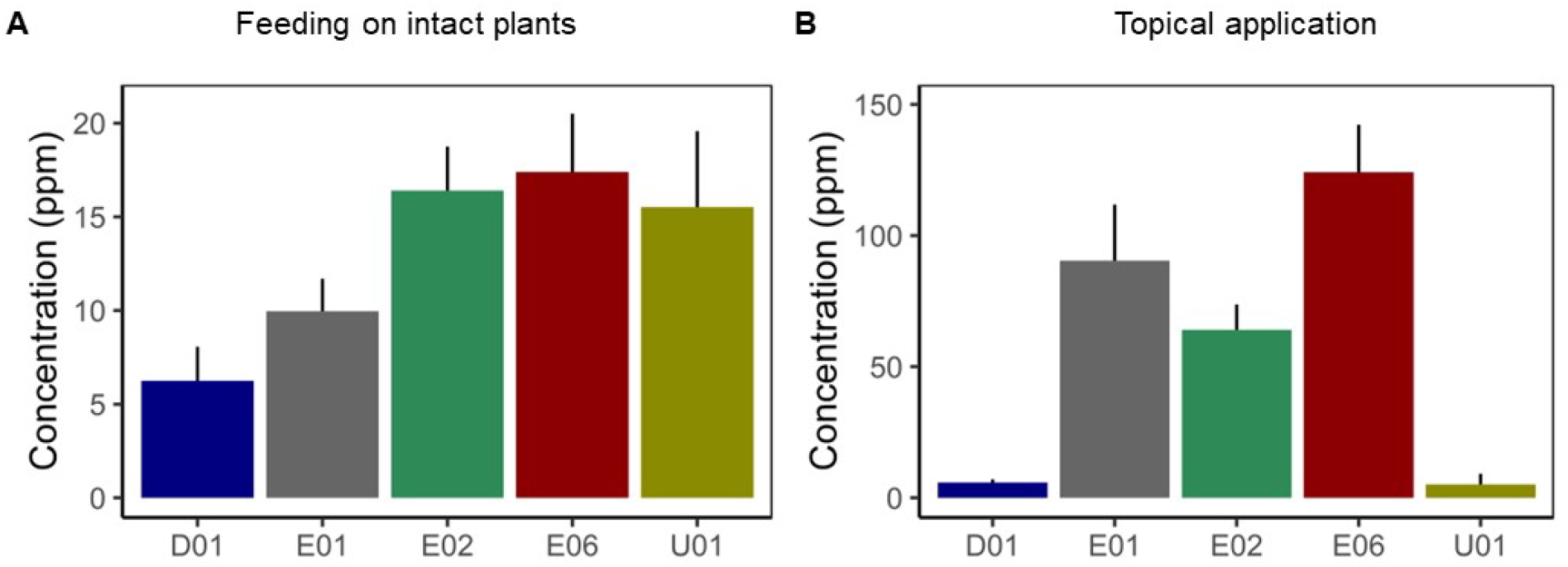
Levels of imidacloprid tolerance differ among five studied Colorado potato beetle strains. LC50 values for two-day-old larvae of the five strains when, **A**. they are allowed to feed on insecticide treated plants, and **B**. when insecticide-water solution is applied topically. The concentration of imidacloprid in the insecticide-water solution is on the Y-axis in ppm. The colors represent the five different strains as indicated on the X-axis.

### B. Behavioral avoidance in larvae

To examine the behavioral avoidance at larval stage, we performed choice assays in Petri dishes using the D01 and E06 strains. The freshly hatched naïve larvae of both strains did not show any preference between control and insecticide-treated plants, suggesting no innate preference of CPB larvae. However, after one day of pre-feeding on potato plants, either with or without the insecticide treatment, the larvae of both strains showed significant preference for control plants (Figure 2).

**Figure 2.**
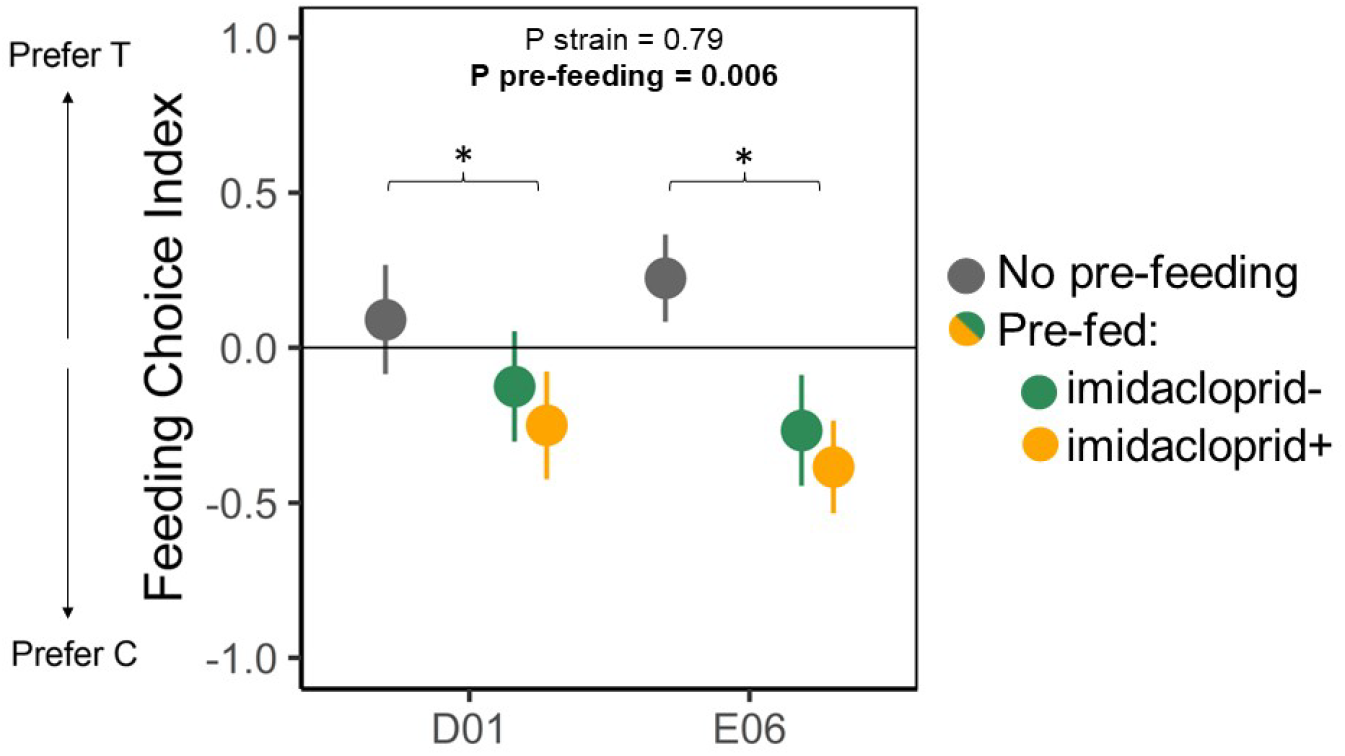
Larvae do not show any innate feeding preference, but pre-feeding affects choice. Choice index is plotted for both the susceptible D01 strain and the resistant E06 strain. The first choice was recorded within 4h after the larvae were introduced into the Petri dishes. The solid horizontal line corresponds to a choice index of 0, meaning that the number of that chose either C or T is equal. Bars represent standard error of the mean. The colors represent the pre-feeding treatment [Grey= no pre-feeding, Green= pre-fed with control leaves (imidacloprid-), and Orange= pre-fed with insecticide-treated leaves (imidacloprid+)]. The horizontal brackets and asterisks represent the significant difference of the pre-feeding treatments (both Control and Treated) from the no pre-feeding treatment.

During the observation period of 4h, no larvae from either strain switched positions from the first chosen disk to the other in any of the treatments. Additionally, first choices were not influenced by the lethal effects of the insecticide since the larvae that died during the assay died before making a choice. To be specific, in the test with naïve larvae, 7 out of 60 D01 larvae and 1 out of 60 E06 larvae died before making a choice. In the assay with pre-fed larvae, for both treatments (pre-fed on control plants and pre-fed on insecticide-treated plants), 1 D01 larva each died before making a choice. In the case of E06, 1 larva pre-fed on control plants died and all larvae pre-fed on treated plants were alive.

### C. Behavioral avoidance in adults

We further measured behavioral avoidance of the two strains at the adult stage using choice assays. When control and insecticide-treated plants were present in the same cage, the first choice (within 30 min) of both the D01 and E06 strains appears to be random (P=0.81, Figure 3A). No differences were found between males and females (P=0.38). Detailed observations on the presence of adults on control and insecticide-treated plants revealed interesting movements and feeding patterns of the two strains. At the start, for both strains there seem to be an equal number of adults on both of the choices of plants. From day 4, the D01 strain (susceptible strain) gradually moved towards controls plants, while the E06 still did not show clear host preference. The strong preference on control plants in the D01 strain continued until the 7th day (Figure 4A). As the consequence, the D01 strain consumed significantly more from the control plants while the E06 strain showed a tendency in the opposite direction (Figure 3B). From day 7, the number of D01 beetles on either plant is not significantly different. This increased movement towards the treated plants is most likely due to reduced eatable leaf material left on the control plant.

**Figure 3.**
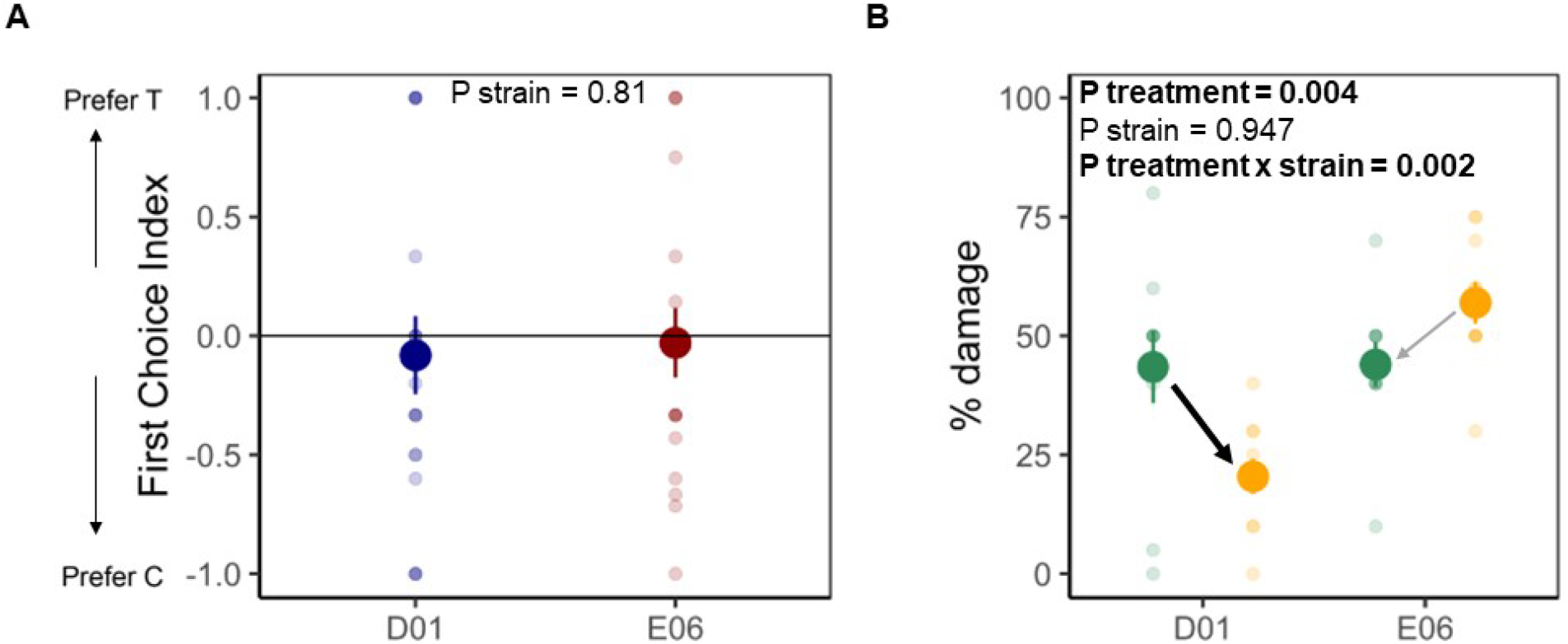
Adults do not show any innate feeding preference, but D01 adults consume more control leaves over time. **A**. First choices of adult beetles within 30min. Choice index graphs showing the preference of beetles in the bioassay with intact plants. The beetles were counted within the first 30min of being introduced into the cages. The solid horizontal line corresponds to a choice index of 0, meaning that the number of beetles on the two choices of plants are equal. The solid dots represent the average number of beetles, and the faded dots show the value for each replicate. Bars represent standard error of the mean. The colors represent the two strains (Blue= D01 and Red= E06). **B**. Percentage leaf damage from intact plants after 10 days. Mean percentage of leaves consumed in the choice assay with adult potato beetles is plotted for both the susceptible D01 strain and the resistant E06 strain as shown on the x-axis. The solid dots represent the average consumption, and the faded dots show the value for each replicate. Bars represent standard error of the mean. The colours represent the two choices of plants (Green= Control and Orange= Treated). The arrows indicate the effect of the interaction of treatment and strain. The thick black arrow indicates the significant preference, and the thin gray arrow indicates the non-significant preference.

**Figure 4.**
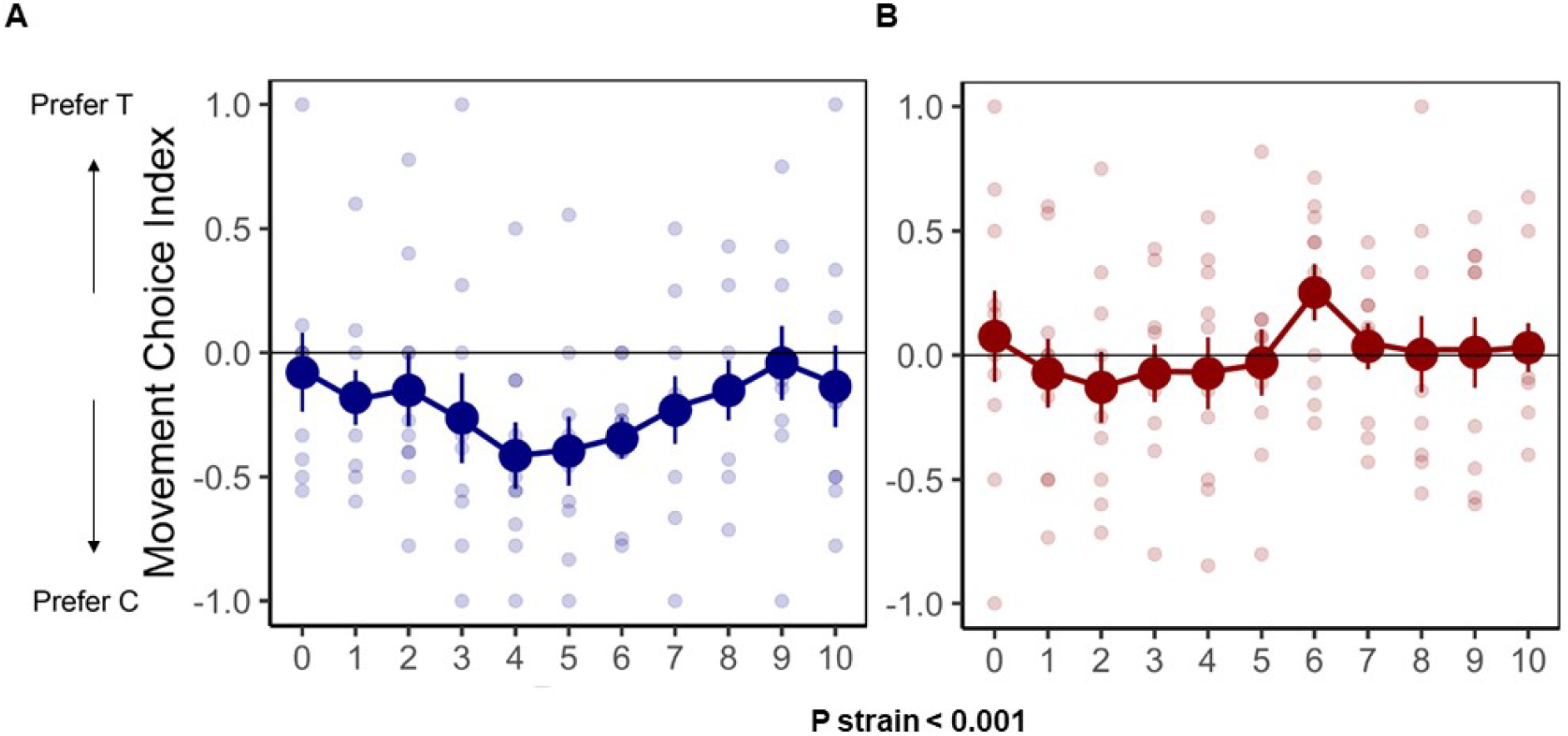
A susceptible strain gradually occupies control plants. The average number of beetles found on insecticide-treated (T) or control (C) plants every day for 10 days is plotted here in the form of a movement choice index. The solid dots represent the average number of beetles, and the faded dots show the value for each replicate. The line segment connecting the solid dots show the movement pattern. Bars represent standard error of the mean. The colors represent the two strains (Blue= D01 and Red= E06).

Based on the eggs collected on each host plant, we estimated the oviposition preference for female adults. Overall, the E06 strain laid about 3 times more eggs in total than the D01 strain (Figure 6A). While the D01 strain laid 73.4% of the eggs on control plants, no strong preference was found for E06 strain (Figure 6A and 6B). The daily oviposition record showed that the D01 strain started laying eggs two days later than the E06 strain and exclusively laid eggs on control plants in the first two oviposition days (Figure 5A & B).

**Figure 5.**
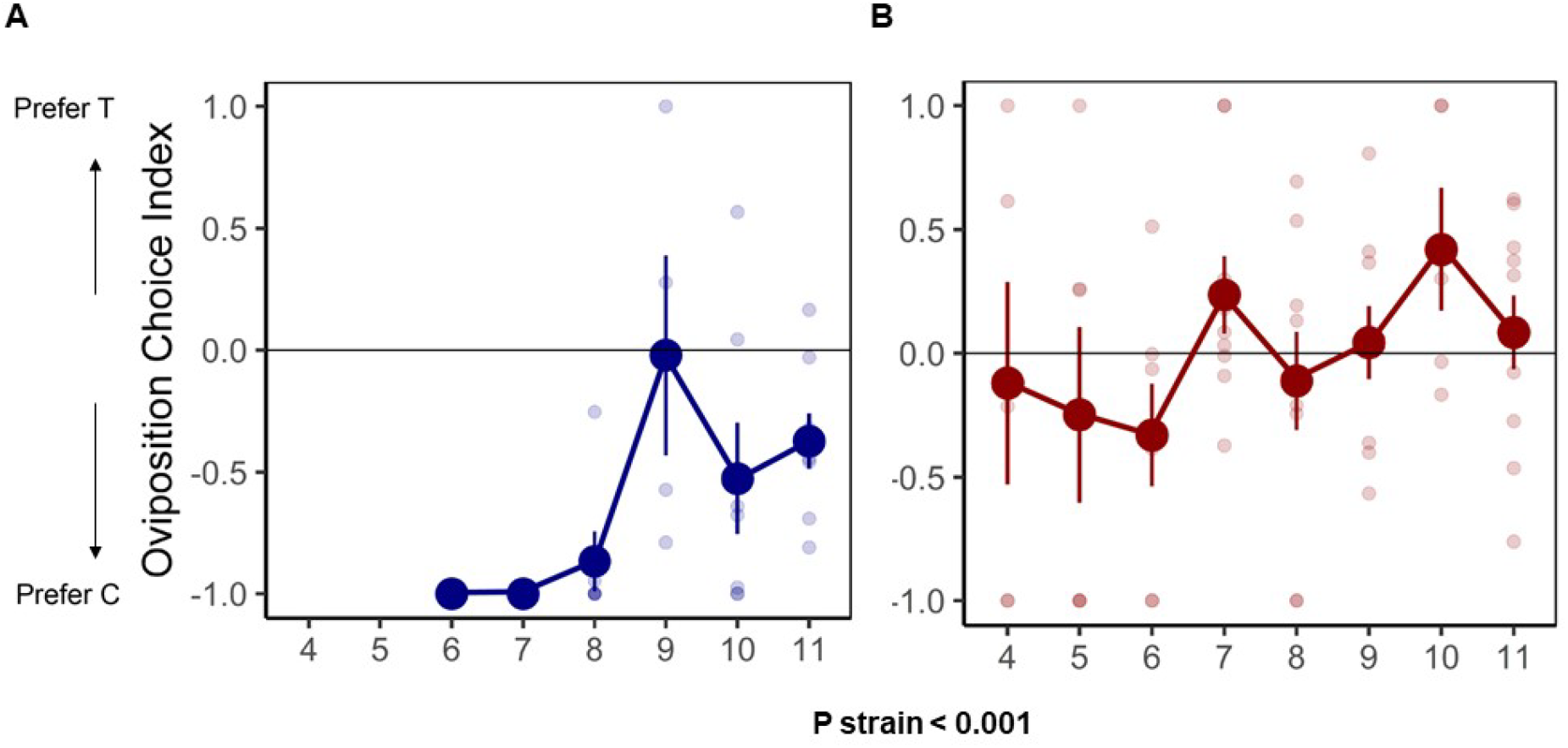
A susceptible strain actively chooses control plants for oviposition. The average number of eggs found on each of the plants every day for the duration of the experiment is plotted here in the form of an oviposition choice index. The solid dots represent the average number of eggs, and the faded dots show the value for each replicate. The line segment connecting the solid dots show the oviposition preference pattern. Bars represent standard error of the mean. The colours represent the two strains (Blue= D01 and Red= E06).

**Figure 6.**
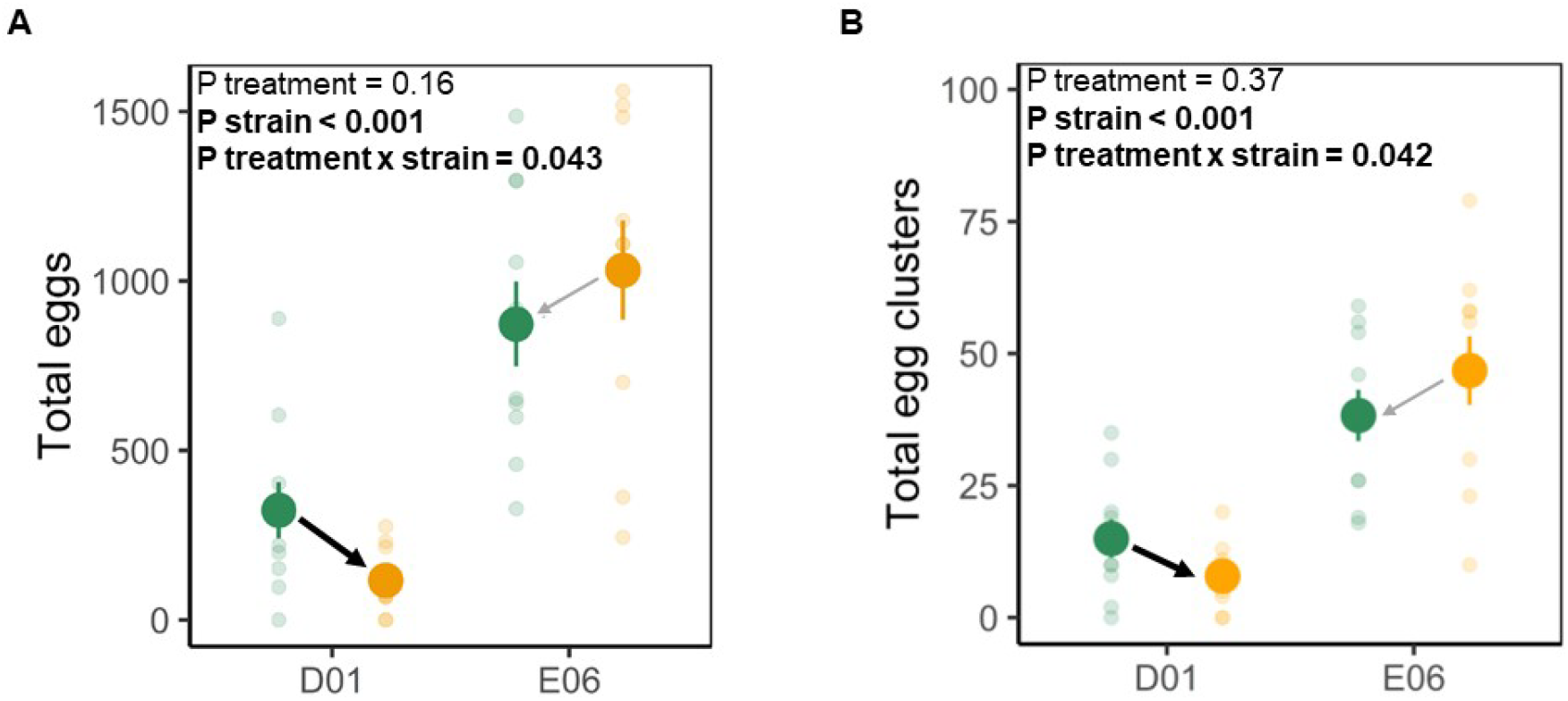
A susceptible strain lays more eggs on control plants than insecticide-treated plants. Total number of eggs and egg clusters per replicate is plotted for both the susceptible D01 strain and the resistant E06 strain as shown on the x-axis. The solid dots represent the average number of eggs, and the faded dots show the value for each replicate. Bars represent standard error of the mean. The colors represent the two choices of plants (Green= Control and Orange= Treated). The arrows indicate the effect of the interaction of treatment and strain. The black arrow indicates the significant preference, and the faded gray arrow indicates the non-significant preference.

## 4. Discussion

Analyzing the levels of behavioral resistance is critical for understanding the evolution of insecticide resistance and host plant adaptation in general. Here, we showed that susceptible CPB adults showed clear behavioral avoidance, both in terms of feeding and oviposition choices, of toxic host plants. Additionally, we showed that these preferences are likely active choices made as a result of experience on insecticide-free and control plants.

We found some similarities and differences between larvae and adults in the behavioral avoidance of imidacloprid. Concerning the similarities, both naïve larvae and adults show no preference when it comes to first choice. However, pre-feeding leads to a preference for control plants. Concerning the differences, the larvae will choose between control or insecticide treated leaf discs and they stay there. That is, the larvae do not actively choose between the hosts. This may suggest that active avoidance behavior probably appears in the later life stages of the beetles. Another difference is that in the case of larvae the strain does not have an effect in the choices.

The movement pattern data shows that the D01 strain prefers to feed on the control plants. For such a preference to manifest, the beetles must detect differences between hosts which can be facilitated primarily by olfactory and/or gustatory cues (29,46). On the one hand, while olfactory cues, such as plant volatiles, play an important role in the host and mate finding behavior of the potato beetles (23,26,47), it is unlikely that they play a significant role here for two reasons. First, there is little evidence that application of imidacloprid via soil affects potato volatiles, although lowered levels of green leaf volatile emission upon spraying imidacloprid was previously reported in tea plants (48). In addition, the distance between control and insecticide treated plants as well as the size of the cages are small, which reduces the likelihood of detecting different volatiles for CPB. Second, first feeding choice is random both in the case of larvae and adults, which could indicate that there are no detectable differences in volatiles between insecticide-free and insecticide-treated plants inside our test cages. On the other hand, CPB are known to have sensitive galeal gustatory cells that respond distinctly to potato leaf alcohols, sugars, amino acids, and their specific combinations (30,32). While the effect of systemic uptake of imidacloprid on the concentration of amino acids and sugars in potato leaves is a possible reason, a more likely mechanism is the detection of insecticide itself via the gustatory system. Further electrophysiological and behavioral studies on the effects of insecticides on gustatory responses are needed to test this hypothesis.

Oviposition choice by the female adults will directly affect the survival of the less motile young larvae of the next generation. We found that the D01 strain laid more eggs on the insecticide-free plants. This could be due to its high susceptibility to the insecticide, that might act as a strong selection pressure in the field and influence traits that affect both feeding and oviposition preferences. Consistently, the resistant strain, E06, did not show a clear preference between the two hosts. While the differences in feeding and oviposition preferences between the two genotypes might be due to the differences of their physiological or biochemical resistances, future studies are needed to demonstrate the underlying associations.

Learning, which influences the host preference in many insects (34–36), likely has contributed to the observed feeding and oviposition avoidance in the D01 strain. This can be seen from the gradual increase of preference to non-toxic plants after 3-4 days in the choice experiment with adults (Fig. 4). The preference disappeared after 7-8 days, likely due to the low availability of plant material and increased competition. The observed learning effects are also associated with oviposition behavior. The D01 strain started laying eggs on day 6 and nearly all eggs were on control plants. Aside from learning, the higher proportion of eggs found on control plants can also be due to the physiological effect of consuming toxic plants, which can reduce or delay the number of eggs laid by the beetles(16). However, we think this is less likely given the correlated changes in their movement (Fig. 4) and oviposition choice patterns (Fig. 5). Future studies that identify the genetic basis underlying the observed oviposition preference will shed further light on how learning contributes to rapid host adaptation in CPB.

## Ethics statement

The experiments were carried out following the 3R Principle put forth by the German Centre for the Protection of Laboratory Animals (Deutsches Zentrum zum Schutz von Versuchstieren, Bf3R: https://www.bf3r.de/). All insects used in the experiments were frozen at −20°C immediately after use.

## Author’s contributions

SX conceived the study and acquired the funding. SX and AE designed the study. RN supplied the five CPB strains and provided valuable information on maintenance and the toxicity assays. AE carried out the experimental rearing and behavioral assays with major aid from AM, DS and LS. AE, AM, DS and LS contributed to data acquisition. MS created the insecticide quantification protocol and performed the analysis. AE wrote the manuscript, with substantial input from PD and SX. PD carried out the statistical analysis. SX and PD supervised the study. All authors gave final approval for publication.

## Data availability

All data used in this study are available along with the R-scripts at https://www.dropbox.com/sh/zpuquedsefrvvdj/AAAKQ39l4Ey8bklGh3guXNasa?dl=0

## Acknowledgements

We thank Holger Schön and David Martin Fernandez for building the insect cages. We are also grateful to Dirk Schmidt and Sascha Ahrens for tips and assistance in growing plants at the greenhouse. Special thanks to Nijat Narimanov for comments on the final manuscript.

## Supplementary Figures

**Supplementary Figure 1.**
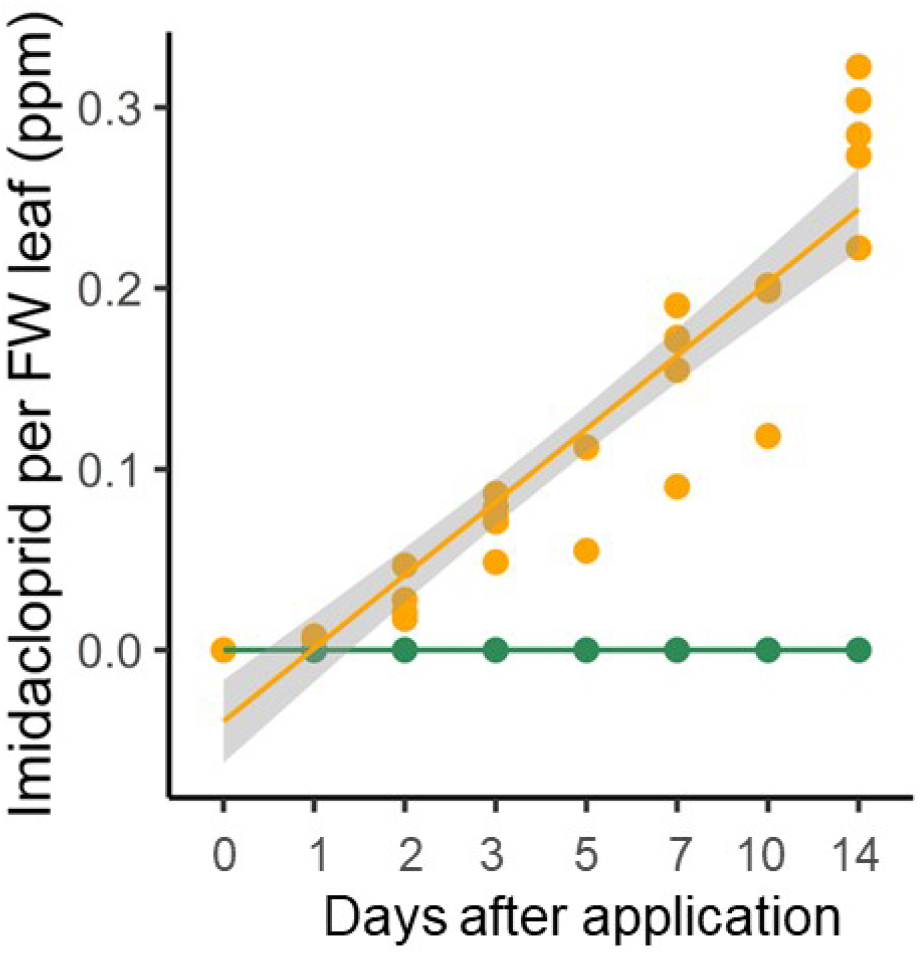
Imidacloprid quantification in potato leaves following soil application. The amount of imidacloprid quantified in the middle leaves (position 3-5 from the top) of 5-week-old potato plants is plotted against the number of days after soil application of 2ppm insecticide-water solution. The colors represent the two treatments of plants (Green refers to control and orange refers to insecticide-treated). The dots represent each replicate (n=3 for control and n=5 for insecticide-treated). The regression lines are denoted by the solid lines and the confidence interval is shown in gray.

## Supplementary Tables

**Supplementary Table 1.** Multi-reaction-monitoring-settings and retention time for LC-MS-based imidacloprid quantification.

|  | RT [min] | Q1 → Q3 [m/z] | Dwell time [ms] | CE | Q1/Q3 Pre Bias [V] |
| --- | --- | --- | --- | --- | --- |
| Imidacloprid | 3.45 | 255,95 175,10 | 97 | -19 | -17 / -17 |
|  |  | 255,95 209,05 | 97 | -17 | -25 / -23 |
| D<sub>4</sub>-Imidacloprid | 3.48 | 259,90 179,20 | 97 | -20 | -17 / -19 |
|  |  | 259,90 213,05 | 97 | -20 | -20 / -21 |

RT: retention time
CE: collision energy
Qualifiers are depicted in grey

**Supplementary Table. 2.** LC50 and LC90 values for 5 strains of potato beetle larvae from the toxicity assay by topical application. The lowest concentrations of imidacloprid in the insecticide-water solution required to kill 50% and 90%, respectively, of the two-day-old potato beetle larvae are reported here alongside standard error (six replicates).

| Strain | LC50 (ppm) |  |  | LC90 (ppm) |  |  |
| --- | --- | --- | --- | --- | --- | --- |
|  | Mean | ±SE | RR | Mean | ±SE | RR |
| D01 | 5.82 | ±1.23 | 1 | 13.46 | ±2.35 | 1 |
| E01 | 90.42 | ±21.42 | 15.54 | 335.36 | ±67.49 | 24.91 |
| E02 | 64.08 | ±9.68 | 11.01 | 123.41 | ±21.52 | 9.17 |
| E06 | 124.16 | ±18.12 | 21.33 | 274.75 | ±43.38 | 20.41 |
| U01 | 5.09 | ±4.10 | 0.87 | 47.24 | ±10.92 | 3.51 |

**Supplementary Table. 3.** LC50 and LC90 values for 5 strains of potato beetle larvae from the toxicity assay using intact plants. The lowest concentrations of imidacloprid in the insecticide-water solution required to kill 50% and 90%, respectively, of the two-day-old potato beetle larvae are reported here alongside standard error (two replicates).

| Strain | LC50 (ppm) |  |  | LC90 (ppm) |  |  |
| --- | --- | --- | --- | --- | --- | --- |
|  | Mean | ±SE | RR | Mean | ±SE | RR |
| D01 | 6.25 | ±1.81 | 1 | 18.88 | ±2.90 | 1 |
| E01 | 9.96 | ±1.74 | 1.59 | 25.52 | ±2.94 | 1.35 |
| E02 | 16.41 | ±2.34 | 2.63 | 41.53 | ±4.69 | 2.20 |
| E06 | 17.40 | ±3.11 | 2.78 | 53.59 | ±6.57 | 2.84 |
| U01 | 15.52 | ±4.06 | 2.48 | 63.75 | ±8.54 | 3.38 |

